# How did SARS-CoV-19 spread in India from Italy, Iran and China? Genetic surveillance of early cases and virus demography

**DOI:** 10.1101/2020.04.26.062406

**Authors:** Mukesh Thakur, Abhishek Singh, Bheem Dutt Joshi, Avijit Ghosh, Sujeet Kumar Singh, Neha Singh, Lalit Kumar Sharma, Kailash Chandra

**Affiliations:** Zoological Survey of India, New Alipore, Kolkata-700053, West Bengal, India; Department of Evolutionary Biology and Environmental Studies, University of Zurich, CH-8057, Zurich, Switzerland

**Keywords:** SARS-Cov-19, phylogenetic analysis, tracing introduction, population demography, Network, substitution rate

## Abstract

SARS-CoV-19 after emerging from Wuhan, drastically devastated all sectors of human life by crushing down the global economy and increased psychological burden on public, government, and healthcare professionals. We manifested by analyzing 35 early coronavirus cases of India, that virus introduction in India, occurred from Italy, Iran and China and population demography apparently revealed a rapid population expansion after the outbreak with a present steady growth. We depicted nucleotide substitutions in structural genes, drove for the adaptive selection and plead for sequencing more genomes to facilitate identification of new emerged mutants, genetic evolution and disease transmission caused by coronavirus.

## Introduction

The Severe Acute Respiratory Syndrome Coronavirus 2 (SARS-CoV-2), emerged at Wuhan city of China in the early December of 2019 [1], resulted out high fatality and incapacitated health systems by causing respiratory illness in human [2]. The virus spread rapidly after the outbreak and took a heavy toll on human life, by causing 187 705 deaths over 213 Countries, areas or territories till dated 25th April 2020 [3]. World Health Organization (WHO) declared SARS-CoV-2 as a global pandemic and issued guidelines to develop cost effective candidate therapeutics for diagnosis of coronavirus. The SARS-CoV-2, being zoonotic in nature, originated from bats [4] and transmitted to human through pangolins as possible intermediate host [1]. Study suggests that East Asian populations plausibly contained different susceptibility to coronavirus due to relatively higher ACE2 expression in tissues, a receptor for SARS-CoV-2 [5]. Due to severe havoc caused by coronavirus, scientists across the globe started analyzing viral genomes to address evolutionary history [6], phylogenetic relationship [7, 8], virus evolution [9] and entry routes in different countries [10, 11]. In this context, the Global Initiative on Sharing Avian Influenza Data (GISAID; https://www.gisaid.org/), founded in 2006, developed a dedicated coronavirus repository in December 2019.

In India, the very first case was reported from a medical student of Thrissur, Kerala having travel history of returning to India from Wuhan on 24th January, 2020. The second and third cases were also reported from Kerala with Wuhan travel history [12]. Eventually, many incidences emerged from different parts of the country with an official record of 19868 active cases, 5803 cured with 824 deaths over 32 States/ Union Territories of India till 26th April 2020, 08:00 GMT+5:30 (https://www.mygov.in/covid-19/?cbps=1). We aimed to investigate the first few possible waves of coronavirus introduced to India, by assigning the early cases to their source of origin and investigate virus demography and adaptive potential by mapping nucleotide substitutions in their structural proteins.

## Materials and methods

### Data mining

We downloaded 35 genome sequences of SARS-CoV-2 available from India, one each of bat and pangolin SARS-related coronavirus, SARSr-CoV (EPI_ISL_402131 and EPI_ISL_410721) from the GISAID database (table S1). These sequences also include the first two cases of SARS-CoV-2 (EPI_ISL_413522 and EPI_ ISL_413523) identified from Kerala. In addition, we included the first Wuhan 1 reference sequence, (henceforth, Wuhan-Hu-1; GenBank ID-MN908947.3), three of the first 16 cases observed in Italy (EPI_ISL_417447, EPI_ISL_417445, EPI_ISL_417446) and a reference genome from Iran (EPI_ISL_424349). All sequences were aligned using MUSCLE v3.8.31, with the Wuhan-Hu-1 [13].

### Genetic diversity, phylogenetic relationship and network analyses

Genetic diversity estimates *i.e.* number of polymorphic sites (P), mean number of pairwise nucleotide differences (K), haplotype (Hd) and nucleotide diversity (π) were calculated using DnaSP v6.12 [14]. Bayesian based phylogenetic analysis was performed using BEAST v1.10.4 [15] with strict clock following HKY substitution model and 10^7^ MCMC simulations. Median-joining network [16] was obtained using NETWORK v4.5.1.0 to study distribution and relationship among the haplotypes with default values for the epsilon parameter (epsilon = 0).

### Demography assessment

Parameters like mismatch distribution test, Harpending’s raggedness index (Rg) and Sum of Squared Deviations (SSD) were calculated to test for demographic expansion using Arlequin v3.5 [17]. Neutrality tests, Tajima’s D [18] and Fu’s Fs [19], to evaluate demographic effects were carried out using DnaSP v6.12 [14]. To date changes in the effective population size over time, we used Coalescent Bayesian Skyline Plot (BSP) analysis as implemented in BEAST v. 2.5 [20] with 8 × 10^^−4^ subs per site per year substitution rate [8, 21]. Markov chains were run for 2.5 × 10^7^ generations and sampled at every 1000 generations, with the first 1000 samples discarded as burn-in. Other parameters were set as default values and output was visualized in the TRACER v. 1.6 [27].

### Mapping nucleotide substitutions in Structural proteins

We mapped mutations occurred in the Spike protein, considered imperative for the adaptive selection [9, 28] using a dedicated, Coronavirus Typing tool of Genome Detective v.1.13 [29], which also maps genetic diversity and mutations with respect to the query sequence. This tool takes in to account Blast and Phylogenetic methods to characterize genomes and corresponding genotypes and identifies both protein and codon mutations in each gene. We classified nucleotide substitutions (dN/dS ratio) in all sequences, with bat RaTG13, Pangolin- SARSr-CoV, SARS and MERS using synonymous nonsynonymous analysis programme, SNAP v.2.1.1 [30]. The dN/dS ratio evaluates the selection pressure, rate of genetic adaptation between structural genes of the diverged species of coronaviruses. If, natural selection promotes and drives for virus mutation by means of fixing non-synonymous change in the protein, the dN/dS ratio is expected to be greater than one, if it suppresses the mutation is expected to be eliminated by purifying selection; the ratio is expected to be less than one.

## Results and discussion

### Genetic polymorphism, phylogeny and network analyses

We obtained 24 haplotypes, 77 polymorphic sites, average nucleotide difference of 10.62 with 0.937 haplotype diversity and 0.00036 nucleotide diversity (table S2). The high haplotype diversity with low nucleotide diversity suggests a recent divergence and high evolutionary potential in coronavirus. Bayesian tree showed the first novel Wuhan-Hu-1 clustered with the first two coronavirus cases reported from Kerala (figure 1a). The phylogenetic tree showed a recent divergence of SARS-CoV-2 from bat SARSr-CoV (divergence time −26.62; 95% CI-24.96-28.33) and Network analysis grouped 24 haplotypes into three major clusters- A, B and C (figure 1b). The cluster A represented by six haplotypes, assigned with three reference genomes of Italy (H_26 to H_28) and cluster B represented by 14 haplotypes assigned with the reference genome of Iran (H_29). Further, the cluster C which represented by only the two haplotypes, H_9 (EPI ISL 413522) and H_10 (EPI ISL 413523), directly linked with the Wuhan-Hu-1, (H_1). Thus, early cases reported from India were genetically assigned in accordance to the patient travel history from Italy, Iran and China (figure 1 a &b). The H_9 was the first coronavirus case reported in India from Thrissur, Kerala and the patient travelled to Kochi, from Kunming via Calcutta on January 24th, and diagnosed positive for SARS-Cov-2 on January 30th 2020 [12]. The H_10 was the third case of Coronavirus in India, with similar travel history [12]. On diagnosis these cases, about 3,400 suspects came in contact with them were put in quarantine to contain the outbreak, however, no further track records available that, how many of them developed coronavirus at later stages. Interestingly, H_12 which had travel history from Iran, genetically assigned with the cluster A, also indicating forward introduction of virus from Italy to Iran. The network tree being monophyletic in nature provides pragmatic evidences to demonstrate that Wuhan, China has been the first source of silent contriver of SARS-Cov-2 to India and thereafter multiple invasion of SARS-CoV-2 occurred in India from Italy, Iran and possibly from many other countries.

**Fig. 1.**
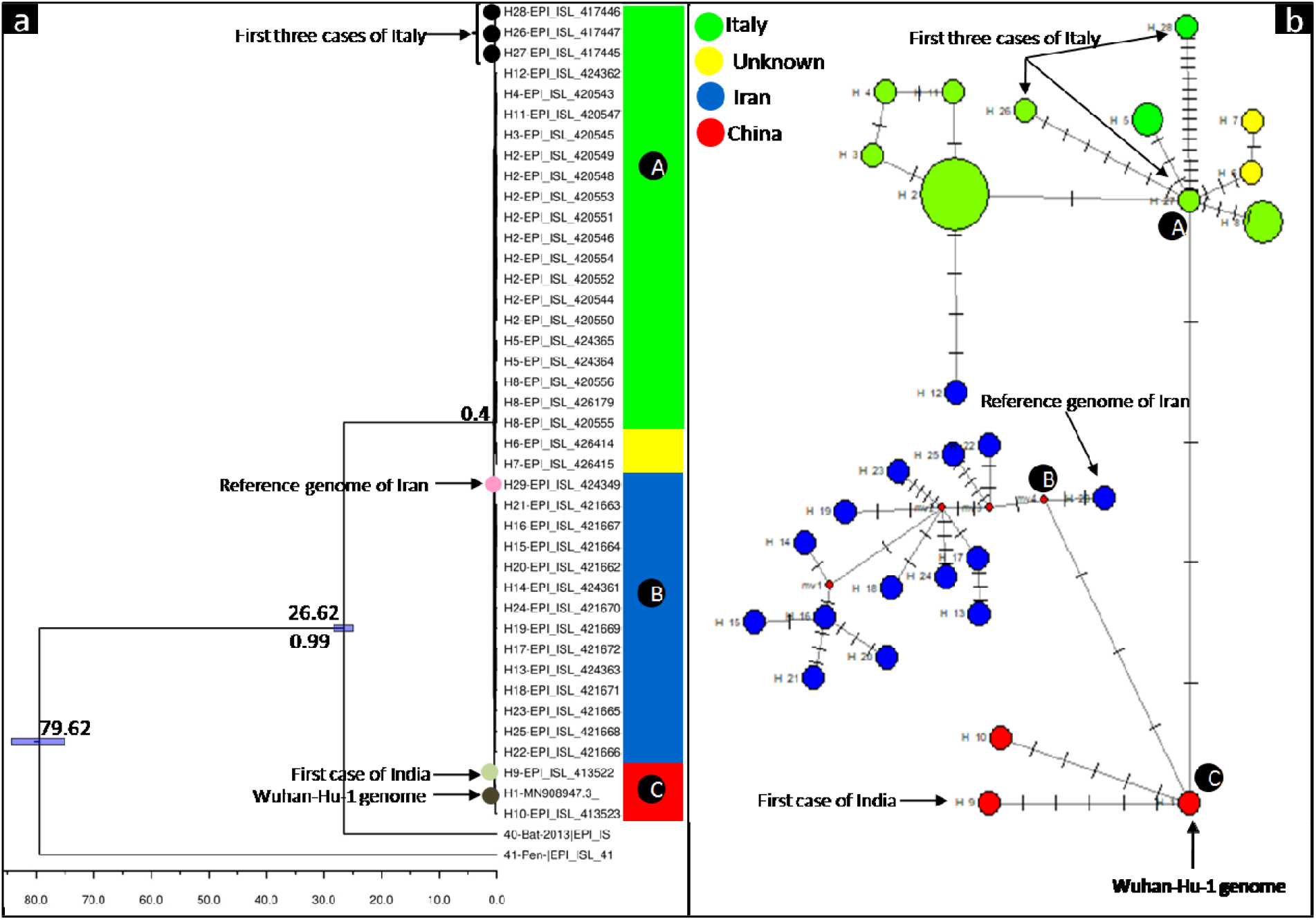
a) Bayesian based phylogenetic relationship among the SARS-CoV-2 genomes. Tree represents 35 SARS-CoV-2 genomes available from India, three reference genomes of Italy, one each reference genome from Iran and China. Bat and pangolin SARSr-CoVs were used as out groups. The scale mentioned on the X axis of the tree is in year and the estimated divergence time of SARS-CoV-2 origin was 0.40 month. **b) Phylogenetic network of SARS-CoV-2 genomes from India.** Haplotype #H_1 in the bright red circle, is the Wuhan-Hu-1 of novel SARS-CoV-2. The other two haplotypes in the red circle, #H_9 and H_10 are the first two cases diagnosed in India which had the immediate travel history from Wuhan, China. Remaining haplotype clusters in green and blue were assigned to their source of origin in accordance to the reference genomes of Italy (haplotypes, H_26 to H_28) and Iran (haplotype,H_29) and travel history of the patients except the haplotype #H_12 which had travel history from Iran but the haplotype matched with Italian clusters. The yellow haplotypes # H_6 and H_7 are foreign travelers but travel history not known. Metadata of all the haplotypes used in the present study is available under the supplementary table S1. The circle areas are in proportion to the number of taxa, and each notch on the links represents a mutated nucleotide position.

### Demography of SARS-CoV-2 in India

We obtained a multimodal pattern of mismatch distribution curve, supporting SARS-CoV-2 is under demographic equilibrium after following a recent expansion (figure 2b). The estimate of Tajima’s D was significant (−1.55132; P <0.5), and also suggested that the population has experienced a recent expansion. The multimodal curve of the mismatch distribution of pairwise genetic differences did not fit a model of sudden expansion in effective population size (SSD 0.02728, *P= 0.270*). Further multimodal mismatch distribution with a non-significant raggedness index (*Rg 0.02728, P= 0.560*), suggested population has maintained a relatively long-term demographic stability after the expansion events (table S2). The BSP revealed a rapid expansion in the effective population size during the last 0.10 to 0.25 year which however became stable and showed a present steady growth (figure 2a).

**Fig. 2.**
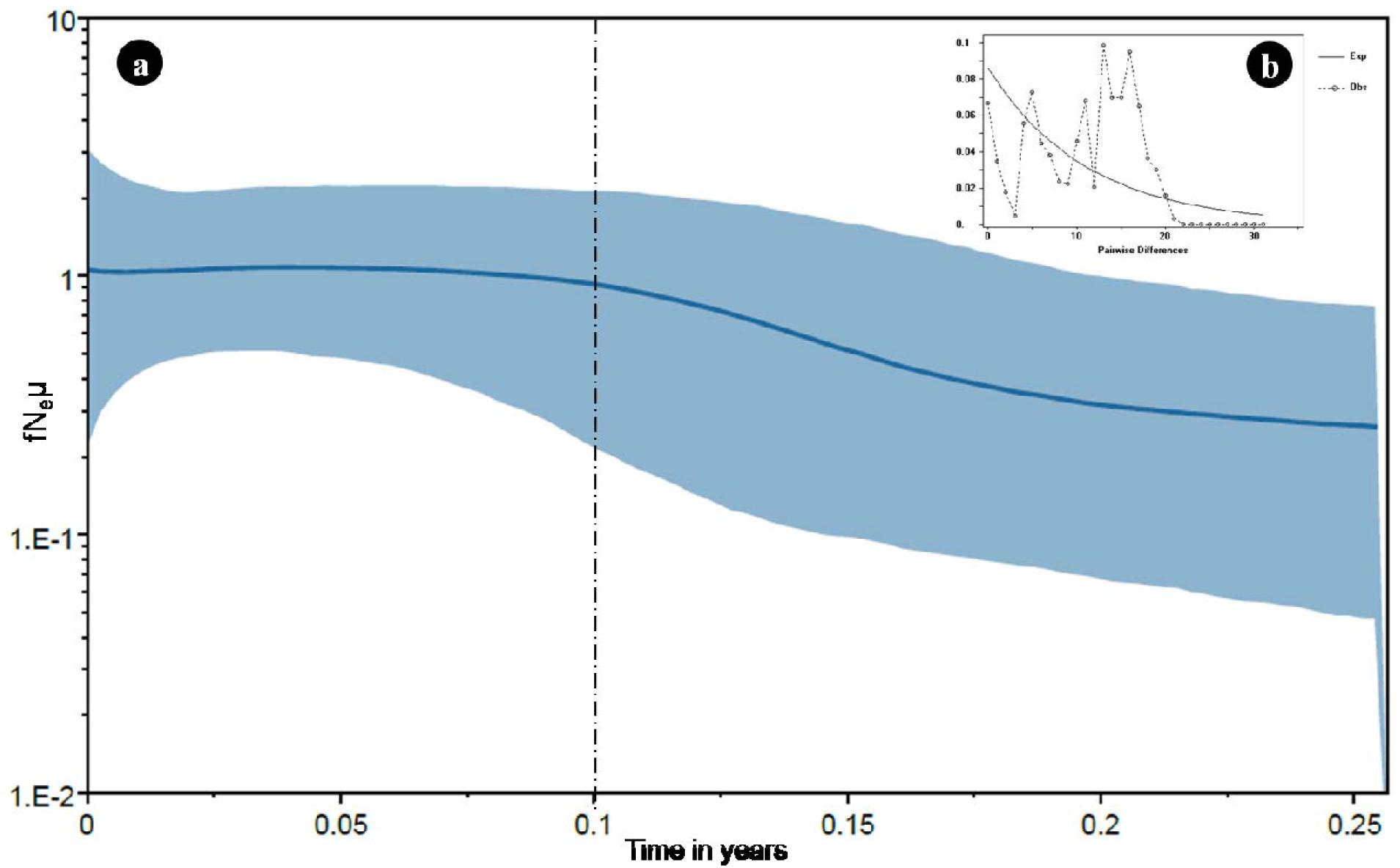
(a) Demographic history of SARS-CoV-2 genomes estimated using Bayesian Skyline Plot. We used a strict molecular clock and a substitution rate of 8 × 10^−4^ /site/year used for SARS-CoV-2 with MCMC run for 2.5 × 10^7^ generations and sampled at every 1000 generations, with the first 2500 samples discarded as burn-in. Other parameters were set as default values and results were visualized in the TRACER v. 1.6 [22]. The time is shown from 0 (present) 0.25 years. Bayesian skyline plot showing an overall stable population size during past last month. The solid line shows the median estimates of Ne μ (Ne = effective population size; μ = generation time), and the blue lines around median estimates show the 95% highest posterior density (HPD) estimate of the historic effective population size. **(b). Mismatch distribution curve of pairwise differences of SARS-CoV-2.**

### Mutation in Spike protein and dN/dS ratio in Structural protein

We found seven unique non-synonymous mutations in the Spike protein to form three clusters, coinciding with Network findings. The cluster A showed a common mutation, D614G (23403A>G) present in all sequences. While, only one genome, EPI_ISL_421671 of cluster B showed a non-synonymous mutation, V622I (23426G>A) in the Spike protein. The cluster C revealed two unique non-synonymous mutations (table S3).

Comparative analysis of dN/dS ratio showed that MERS-CoV was the most diverged and Bat SARSr-CoV was the closest relative to SARS-CoV-2 (table S4). Further, the dN /dS ratio of the envelop (0.318), membrane (0.01) and nucleocapsid (0.198) genes of SARS-CoV-2 with other coronavirus suggested that the changes in the protein sequences was responsible to drive purifying selection [26]. The dN/dS ratio of the Spike gene was the largest (0.663), signifying this region to be under adaptive selection and plausibly helping virus in transmission (supplementary table S4). Interestingly, the dN/dS ratio from a set of conspecific sequences of a single population, may not essentially be greater than one as synonymous mutation often eliminate deleterious mutation to fix the adaptive selection [27]. The mutation of D614G (23403A>G) observed in Spike protein of genomes with travel history to Italy was also reported as a prevalent mutation in the viral isolates from Europe [28], corroborating the source assignment.

We demonstrated, by analyzing 35 high quality genomes that early introduction of the virus in India and its spread was through the cases having travel history from Italy, Iran and China by corroborating the case history, phylogeny and network analysis. The study exhibited, a present steady growth of the virus in India, which indicates virus is still operating and several new corona cases may appear in the near future. Till 26th April 2020, 08:00 GMT+5:30, 26495 coronavirus cases were reported from India and unfortunately only 35 complete genomes of the total database of 12,268 virus sequences were available from India on the growing GISAID database. We plead authorities to kindly issue permits and grant funds to the research institutes/ labs and appeal scientists to populate more genomes of SARS-CoV-2 of the country in the public database. This will provide scientific fraternity a common platform to analyze, collaborate and understand genetic evolution of the virus to emerge into new structural mutants, transmission rate as well as in-time development of diagnostics, therapeutics, vaccines and antivirals.

## Acknowledgements

Authors acknowledge the team GISAID (https://www.gisaid.org/), for providing data access and personalized tool to download and analyze data. The study was compiled within a week, during the lockdown period in India and the first author greatly thank to his wife, Mrs Vertika Sisodiya & daughter, Grace for providing him adequate time and space and allowing him to focus on the present study.

